# The transcription of a single olfactory receptor per neuron is enforced by epigenetic silencing of their enhancers

**DOI:** 10.64898/2025.12.22.695993

**Authors:** Francesco Tabaro, Basilia Acurzio, Aileen Julia Riesle, Monika Mielnicka, Camilla Lyons, Domenico Guarascio, Agnieszka Sadowska, Michela Ascolani, Marina Peralta, Emerald Perlas, Alvaro H. Crevenna, Neil Humphreys, Hiroki Asari, Mathieu Boulard

**Author notes:** These authors contributed equally to this work (Francesco Tabaro, Basilia Acurzio).

## Abstract

The ability to discriminate thousands of odors in our environment requires each olfactory neuron to express a single olfactory receptor from hundreds of available genes. The biochemical mechanism enforcing this monogenic expression remains unknown. We show that deletion of the chromatin protein TRIM66 causes individual olfactory neurons to express multiple receptors at a high level, demonstrating that monogenic expression relies on an epigenetic silencing mechanism. Moreover, TRIM66 is specifically recruited to olfactory receptor gene super-enhancers during neuronal progenitor maturation, thereby silencing nearby olfactory receptor genes. Loss of monogenic expression disrupted axonal projections to the olfactory bulb, resulting in an aberrant topographic map and impaired social odor discrimination and reproductive behaviors. These findings uncover the chromatin-based silencing of super-enhancers as the mechanism underlying the organization of the mammalian olfactory system.

## INTRODUCTION

Animals rely on their sense of smell for their survival, olfaction being essential to find mates and food, to nurse progeny, and to avoid predators. Higher vertebrates perceive their chemical environment through the detection of volatile molecules in the nasal cavity by specialized bipolar neurons called olfactory sensory neurons (OSNs). At the molecular level, the detection of odorant molecules is mediated by G-coupled proteins with seven transmembrane domains termed olfactory receptors (ORs) that are located on the cilia of OSN dendrites ^1^. A wide olfactory receptive field is afforded by an extremely high number of ORs. Rodents, for instance, possess as many as 1,000 and humans approximately 400 ORs ^2^. The genes that encode ORs are organized in genomic clusters of various sizes found on most chromosomes. Their transcription is tightly regulated in a way that individual OSNs express only one OR gene in a mono-allelic fashion, selected in a largely stochastic manner ^3^, a process that occurs during the maturation of OSN precursors ^4,5^. The exclusive transcription of a single OR allele from a large gene repertoire generates extensive cellular diversity without changes in the DNA sequence ^6,7^. Hence, the murine main olfactory epithelium (MOE) is composed of over 1000 types of OSNs distinguished by the OR they express. The “*one neuron–one receptor”* rule is fundamental for olfactory perception because the identity of the chosen OR confers two key properties to the neuron: its ligand specificity and its axon targeting to one of two possible discrete glomeruli in the olfactory bulb (OB) ^8^.

The olfactory epithelium is continuously regenerated during the life of the organism from stem cells called globose basal cells (GBC) that are produced by the differentiation of quiescent horizontal basal cells located in the basal layer of the neuroepithelium ^5^. Intriguingly, the differentiation of GBCs into immediate neuronal precursors (INP) is accompanied by the low-level expression of multiple OR genes ^4,5,9^. Subsequent differentiation of INPs into immature OSNs (iOSNs) induces a transcriptional switch from polygenic to monogenic OR transcription ^5^. Hence, each iOSN expresses a single OR whose expression increases after terminal differentiation into mature OSN (mOSN).

While the molecular mechanism that controls the polygenic-to-monogenic transcriptional switch remains enigmatic, progress has been made in understanding the regulatory DNA sequences and trans-acting factors that drive transcription of OR genes. A major finding was the discovery that a network of approximately 100 enhancers known as the Greek Islands enhancers (GI-enhancers) is essential for OR transcription ^10–12^. GI-enhancers are characterized by open chromatin in mOSNs and their occupancy by the transcription factors EBF1 and LHX2 ^11,12^ which are crucial for OR gene transcription ^11,13^. Each OR gene cluster contains at least one GI-enhancer ^12^, which is required for the transcription of the closest OR genes located in *cis* ^10,14,15^. However, the *cis-*located GI-enhancer is necessary but not sufficient to activate OR transcription, and the *trans*-activity of other GI-enhancers is also required ^13^. The aggregation of GI-enhancers in *trans* occurs during the maturation of OSNs and is promoted by the LIM-domain-binding protein LDB1, which forms a homodimer and binds to LHX2-EBF1 ^13^. Thus, OR gene activation relies on a synergistic combination of *cis-* and *trans*-acting GI-enhancers. However, these findings do not shed light on how OSNs promote the exclusive expression of only a single OR. Here, we report that a chromatin factor named Tripartite Motif-Containing Protein 66 (TRIM66, also known as TIF1-delta) enforces the monogenic transcription of OR genes.

## RESULTS

### TRIM66 is a constituent of olfactory sensory neuron chromatin

TRIM66 was first reported as a male germ cell-specific chromatin factor ^16^. While we found no spermiogenesis defect in male mice homozygous for a loss-of-function *Trim66* allele, homozygous males sired fewer litters than wild types (WT) when bred with WT females for the same period of time (Figure 1A,B). Reassessment of *Trim66* expression across tissue types revealed a previously unappreciated expression in the embryonic nasal epithelium (Figure S1A). Single-cell transcriptomic profiling of the adult main olfactory epithelium (MOE) characterized the dynamics of *Trim66* expression during OSN maturation (Figure S1B,C). To detect TRIM66 protein, we created an epitope-tagged *Trim66* allele by insertion of a haemagglutinin epitope (HA) at the N-terminus of endogenous TRIM66. Immunostaining for TRIM66 in MOE showed specific expression in olfactory neurons (Figure S1D). Co-staining with cell-type markers documented TRIM66’s subnuclear localization in INP, iOSNs and mOSNs. Expression of TRIM66 was not detectable by immunostaining in INPs (Figure S1E), and first became apparent in iOSN where it formed numerous foci throughout the nucleus (Figure S1F). TRIM66 immunostaining pattern changed in mOSN where it showed fewer foci arranged around the chromocenter (Figure 1C). Collectively, these data identified TRIM66 as a constituent of iOSN and mOSN chromatin with a developmental expression coinciding with the onset of OR transcriptional singularity.

**Figure 1:**
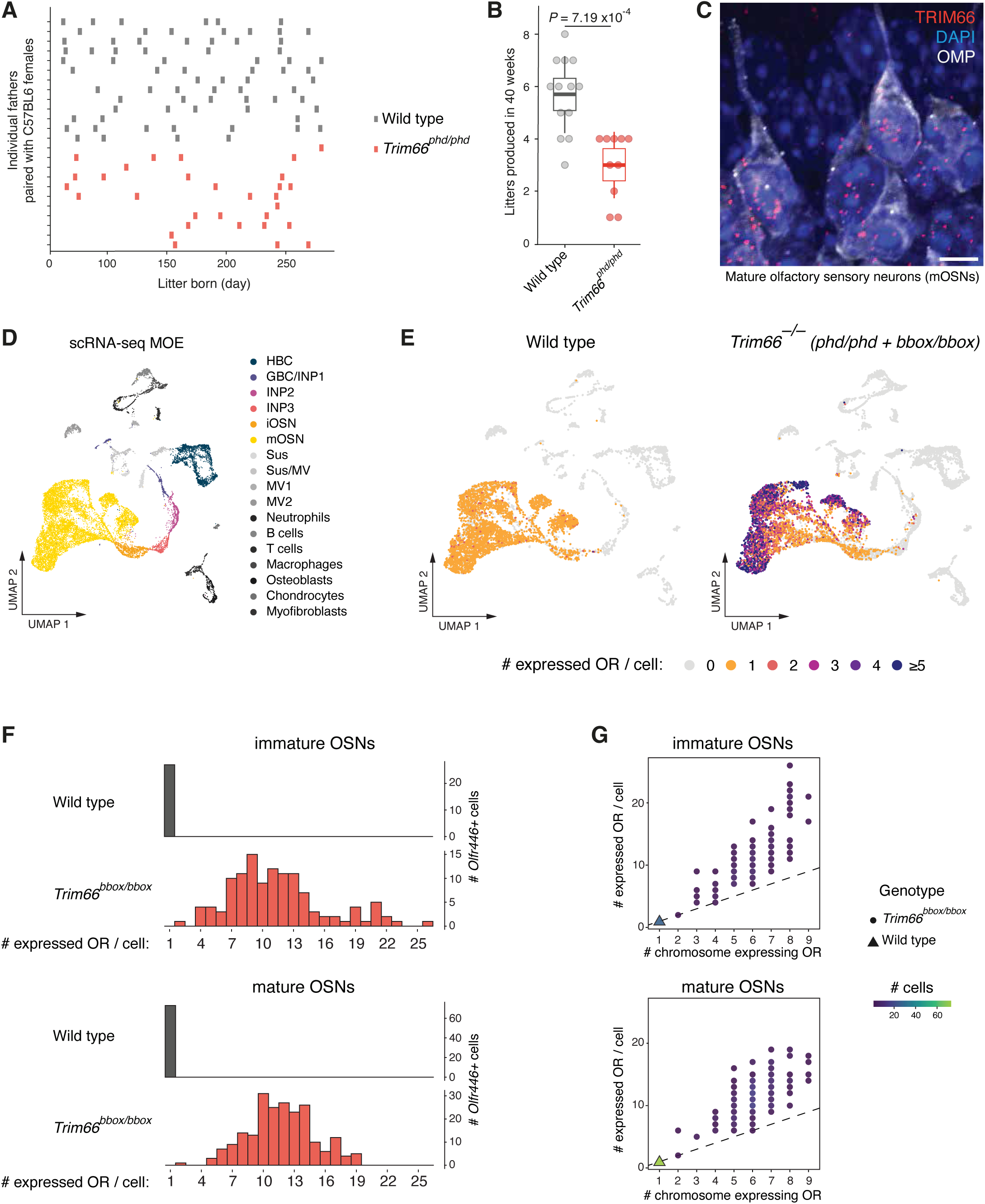
Widespread OR multigenic expression in *Trim66*-deficient iOSNs and mOSNs. (A) Frequency of litter production by *Trim66^phd/phd^* males (n=10) and WT males (n=13) paired with C57BL6 females over a period of 40 weeks. Each row represents an individual male of the indicated genotype, each bar represents a litter. (B) Number of litters produced by *Trim66^phd/phd^* males (n=10) and WT males (n=13), paired with C57BL6 females over a period of 40 weeks. Horizontal lines show median values, boxes represent the 25^th^ to 75^th^ interquartile range, whiskers extend from the 75^th^ quartile + 1.5 * interquartile range and the 25^th^ quartile - 1.5 * interquartile range. Each dot represents a litter size. Significance calculated with Mann-Whitney-Wilcoxon test. (C) Confocal image of a section of MOE showing TRIM66 expression in mOSN. Endogenous TRIM66 fused to the HA epitope tag was detected with anti-HA antibodies (red). OMP staining (white) marks mOSN. DNA staining with DAPI is shown in blue. Scale, 5 µm. (D) Uniform manifold approximation and projection (UMAP) visualization of MOE 10x single-cell RNA-sequencing (scRNA-seq) integrated dataset colored by the assigned cell type. (E) UMAP visualization of MOE 10x scRNA-seq colored by the number of OR genes expressed per cell. Integrated samples from two WT controls (left panel), *Trim66^phd/phd^* and *Trim66^bbox/bbox^*(right panel). (F) Histogram representing the number of OR genes expressed in individual *Olfr446*+ iOSNs (top two panels) and mOSNs (bottom two panels). The genotype is indicated (Wild type, *_Trim66bbox/bbox_*_)._ (G) Dot plot showing the number of chromosomes involved in OR gene co-expression per *Olfr446*+ neuron. Dot color indicates the number of cells, and dot shape denotes the genotype. The dashed line represents the 1:1 ratio. Top: iOSNs; bottom: mOSNs.

### *Trim66* removal disrupts the “one neuron-one receptor” rule

To investigate the functional role of TRIM66, we disrupted the murine *Trim66* gene by insertion of a premature stop codon in exon 15, termed *Trim66^phd^* (Figure S2A) ^17^. Differential gene expression in the whole adult MOE was initially quantified by bulk RNA-seq. Almost all OR genes (n=1,111) were deregulated in homozygous mutant MOE (76 upregulated and 875 downregulated, adjusted *P*-value < 0.01, Wald t-test, log2-fold change > 0.5, Figure S2B). In contrast, other genes were largely unaffected by *Trim66* removal (Figure S2B, right panel). Thus, TRIM66 plays a specialized function in the regulation of OR gene transcription; however, its role is distinct from OR gene trans-activators, like *Lhx2* and *Ldb1,* whose disruption caused the downregulation of nearly all OR genes and no upregulation of OR genes ^11,13^. We hypothesized that the upregulation of OR genes in *Trim66*-deficient MOE could be a consequence of the co-expression of multiple ORs within individual OSNs (loss of singularity). We tested this possibility by quantitating the number of OR mRNAs in individual mOSNs using droplet-based single-cell RNA-seq (scRNA-seq; 10x Genomics). To assess replicability, we analyzed two independent *Trim66* loss-of-function alleles: *Trim66^phd^* as previously described ^17^, and *Trim66^bbox^,* which we created by inserting a stop codon in exon 4 (Figure S2A). Both *Trim66^phd^* and *Trim66^bbo^*^x^ mutations are null, as demonstrated by the absence of detectable TRIM66 protein in the MOE of homozygous and heterozygous compound animals (Figure S2C). Using previously identified marker genes (Table S1), we first clustered cells by cell type (Figure 1D)^5^ and then quantitated OR transcripts per single cell (normalized read count > 5). As expected, only one OR mRNA was detected per WT mOSNs (Figure 1E, left panel). In contrast, TRIM66 deficiency resulted in the co-expression of a variable number of ORs per mOSN ranging from 1 to 11 (median=3), with the vast majority of single mOSNs expressing more than 1 OR gene (Figure 1E, right panel). The widespread derepression of many OR genes per neuron was reproducible between the two *Trim66*-null alleles (Figure S3A-C). Sixty-four percent of the bulk RNA-seq upregulated OR genes (n=49) were among the co-expressed ORs identified by scRNA-seq, supporting the conclusion that OR upregulation at the bulk level was a reflection of multigenic OR expression in single mOSNs (Figure S3D). RNA *in situ* hybridization confirmed the frequent co-expression of two distinct OR genes in mOSNs lacking TRIM66 (Figure S3E). Altogether, these results show that TRIM66 is required for the monogenic expression of OR genes.

### TRIM66 mediates the developmental switch from multigenic to monogenic OR expression

We next explored the developmental origin of OR gene co-expression by profiling the transcriptome at the two key stages of maturation where TRIM66 is normally present, namely iOSNs and mOSNs. To quantify the transcription of OR genes with a higher sensitivity and accuracy, we used a single-cell technique with full-length transcript coverage (Smart-seq2), and we increased the sequencing coverage by focusing on a subpopulation of neurons expressing a particular OR gene. We chose *Olfr446* that is upregulated in *Trim66*-deficient mOSNs (Figure S4A) ^18^. To isolate *Olfr446*-expressing neurons, we created a reporter allele, termed *Olfr446^T2i-TAU-GFP^*, by insertion of a cassette consisting of a self-cleaving T2A tag followed by a TAU-GFP, allowing the visualization of their axons. We computationally separated iOSN (WT n=29; *Trim66^bbox/bbox^* n=120) from mOSN (WT n=73; *Trim66^bbox/bbox^* n=216) based on known marker genes (Table S1)(Figure S4B-D). While WT iOSNs only show monogenic OR expression, *Trim66*-disrupted iOSNs expressed multiple ORs (between 2 and 26, median=11) (Figure 1F, top panel). A similar phenotype is observed in mOSNs, where WT cells express a single OR gene and *Trim66*-deficient cells express between 2 and 19 ORs (median=12) (Figure 1F, bottom panel). The loss of the monogenic transcription was fully penetrant, as all *Trim66*-mutant neurons analyzed (n=336) expressed more than two OR genes (Figure 1F). The totality of the OR mRNA detected in *Trim66*-mutant iOSNs (n=65) were also expressed in *Trim66*-mutant mOSNs, indicating that OR multigenic expression in mOSN is likely a developmental legacy of a disruption of the singular OR transcription in iOSNs. A total of 85 ORs were detected in our population of single neurons, including *Olfr446*; however, it was not always the most expressed OR gene, and no particular pattern emerged in the ranking by expression level (Figure S4E). Next, we asked whether the co-expression of multiple OR genes per neuron impacts their level of expression. Focusing on the most highly expressed OR gene, both *Trim66*-mutant iOSNs and mOSNs exhibited increased expression compared to the singly expressed OR in WT (iOSN: *P*=3.353e^−10^; mOSN: *P*=0.0002082, Kolmogorov-Smirnov test). Within a single neuron, the co-expressed ORs had a similar expression level (median normalized expression value = 58.82; mean variance per cell: 3.44) (Figure S4F). Finally, we examined the genetic linkage of the co-expressed OR genes within a single cell and found that co-expressed OR genes in *Trim66*-mutant neurons were overall located on different chromosomes (Figure 1G). Considering previous evidence that *cis*-acting GI-enhancers are required for the transcription of nearby genes ^10,14,15^, this result indicates that there could be more than one *cis*-acting GI-enhancer driving transcription in *Trim66*-deficient olfactory neurons.

In summary, these data show that removal of TRIM66 results in the aberrant transcription of multiple OR genes from different chromosomes in iOSN, which is the developmental stage where all but one OR gene should be silenced ^4,5,9^. These results, taken together with TRIM66’s onset of expression in iOSN (Figure S1E,F) argues that TRIM66 acts as a molecular switch that enforces the transition from polygenic to monogenic OR transcription during the transition from precursor (INP) to iOSN.

### TRIM66 occupies OR enhancers

We next investigated the genomic occupancy of TRIM66 by performing CUT&RUN on nuclei isolated from adult MOEs. The regions bound by TRIM66 were limited to approximately 50 loci, corresponding to GI-enhancers (Figure 2A). TRIM66 occupied approximately half of the 103 known GI-enhancers (Figure 2B). This result raises the possibility that there might be TRIM66-dependent and TRIM66-independent GI-enhancers with different regulatory functions. This finding contrasts with the known trans-activators of OR genes LHX2, EBF1 and LDB1 that occupy nearly all GI-enhancers (Figure 2B). It is conceivable that the binding of TRIM66 on one hand and that of LHX2/EBF1/LDB1 on the other hand to GI-enhancers could be mutually exclusive. It is noteworthy that TRIM66 was only present at GI-enhancers and that there was no detectable TRIM66 association with OR gene promoters (Figure 2C).

**Figure 2:**
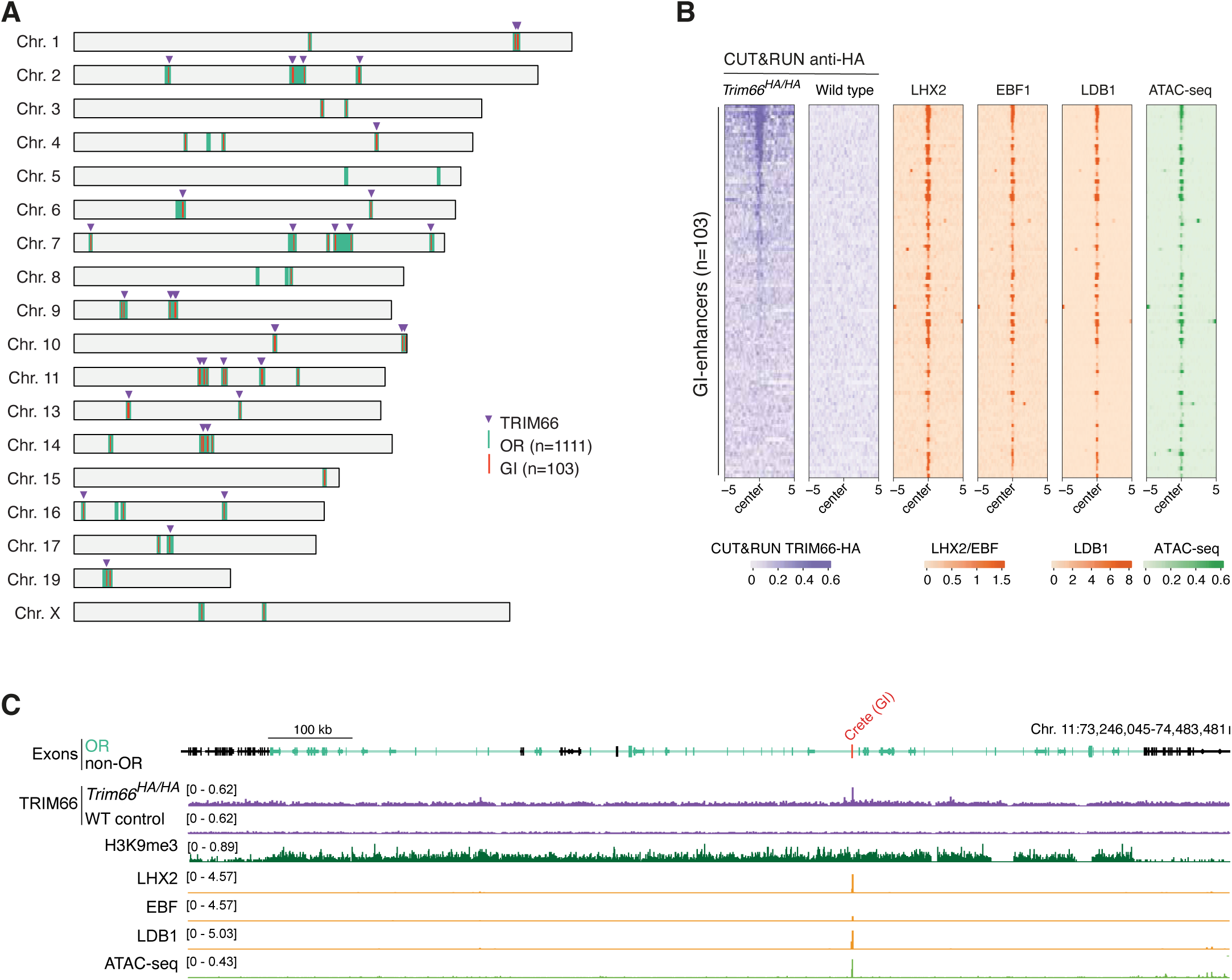
TRIM66 exclusively binds to GI-enhancers. (A) Position of OR genes cluster (n=68) and GI-enhancers (n=103) on the murine chromosomes. The GI-enhancers occupied by TRIM66 (n=50) are indicated with the red arrows. (B) Chromatin profiling at GI-enhancers (n=103). Left panels: TRIM66 occupancy measured by CUT&RUN with anti-HA antibodies in *Trim66^HA/HA^* and WT (no epitope control) MOEs. Middle panels: LHX2, EBF1, LDB1 and ATAC-seq profiling in the isolated population of mOSNs. Each row of the heat map shows 10 kb centered on the GI-enhancer. Heatmaps are ordered by descending TRIM66 CUT&RUN signal intensity. LHX2, EBF (ChIP-seq, mOSNs dataset: GSE93570); LDB1 (ChIP-seq, mOSNs dataset: GSE112153); ATAC-seq (mOSN dataset: GSE93570). (C) Genomic visualization of the OR cluster on chromosome 11 containing the GI-enhancer “Crete” (Chr. 11:73,246,045–74,483,481). OR genes are highlighted in light green, non-OR genes in black. The screenshot displays TRIM66 occupancy tracks in *Trim66^HA/HA^* and WT samples, ChIP-seq for H3K9me3, LHX2, EBF, LDB1, and ATAC-seq.

### TRIM66 prevents promiscuous OR expression in the genomic cluster in *cis*

The physical proximity of GI-enhancers located on different chromosomes in the nuclei of mOSNs is essential for OR gene’ transcription ^13^. However, it was reported that the propensity of different GI-enhancers to engage in *trans*-contacts varies greatly ^10,13^. The variation in *trans*-contact frequency between GI-enhancers is currently not understood and likely reflects variable combinations of trans-interactions between GI-enhancers in the heterogeneous population of mOSN. Using public Hi-C data from isolated mOSNs, we computed the *trans* proximity score for all known GI-enhancers and asked whether these differences might be correlated with TRIM66 occupancy. We found that TRIM66-bound GI-enhancers tended to be located at the center of the GI-enhancer *trans*-interaction network (Figure 3A) and displayed a higher probability of being in physical proximity as compared to GI-enhancers with lower TRIM66 binding (*P*=4.99e^−5^, Wilcox-Mann-Withney U-test) (Figure 3B). Hence, TRIM66 binds to the subset of GI-enhancers that makes the highest number of *trans*-contacts. This result appears paradoxical because *trans*-contacts between GI-enhancers promote OR transcription ^13^, while the data shown in Figure 1 support a repressive role for TRIM66. However, TRIM66 could bind to its target GI-enhancers in their inactive state, which were shown to make *trans*-contact and aggregate in inactive GI-hubs ^12,19^. The requirement of the proximal GI-enhancer in *cis* for transcription raised the possibility that TRIM66 binding could suppress their *cis-*acting function ^10,14^. To test this hypothesis, we compared the number of OR genes that are monogenically or polygenically expressed in *cis* proximity to GI-enhancers in *Trim66*-deficient mOSNs, grouping GI-enhancers by their occupancy by TRIM66 in the WT (Figure 3C, low TRIM66 binding on the left, high TRIM66 binding on the right). This analysis revealed that OR genes located in *cis* of a GI-enhancer normally occupied by TRIM66 are more likely derepressed in *Trim66*-deficient mOSNs than the OR genes in proximity of GI-enhancers with low TRIM66 occupancy (P= 7.56e^−5^, *t*-test, Figure 3D). This result supports a role for TRIM66 in repressing the OR genes located in the *cis* genomic cluster. Given the multiple chromosomal origins of the co-expressed OR transcripts after TRIM66 removal and the higher responsiveness of OR genes located in *cis*, we conclude that TRIM66 binding to an enhancer represses its *cis*-acting function.

**Figure 3:**
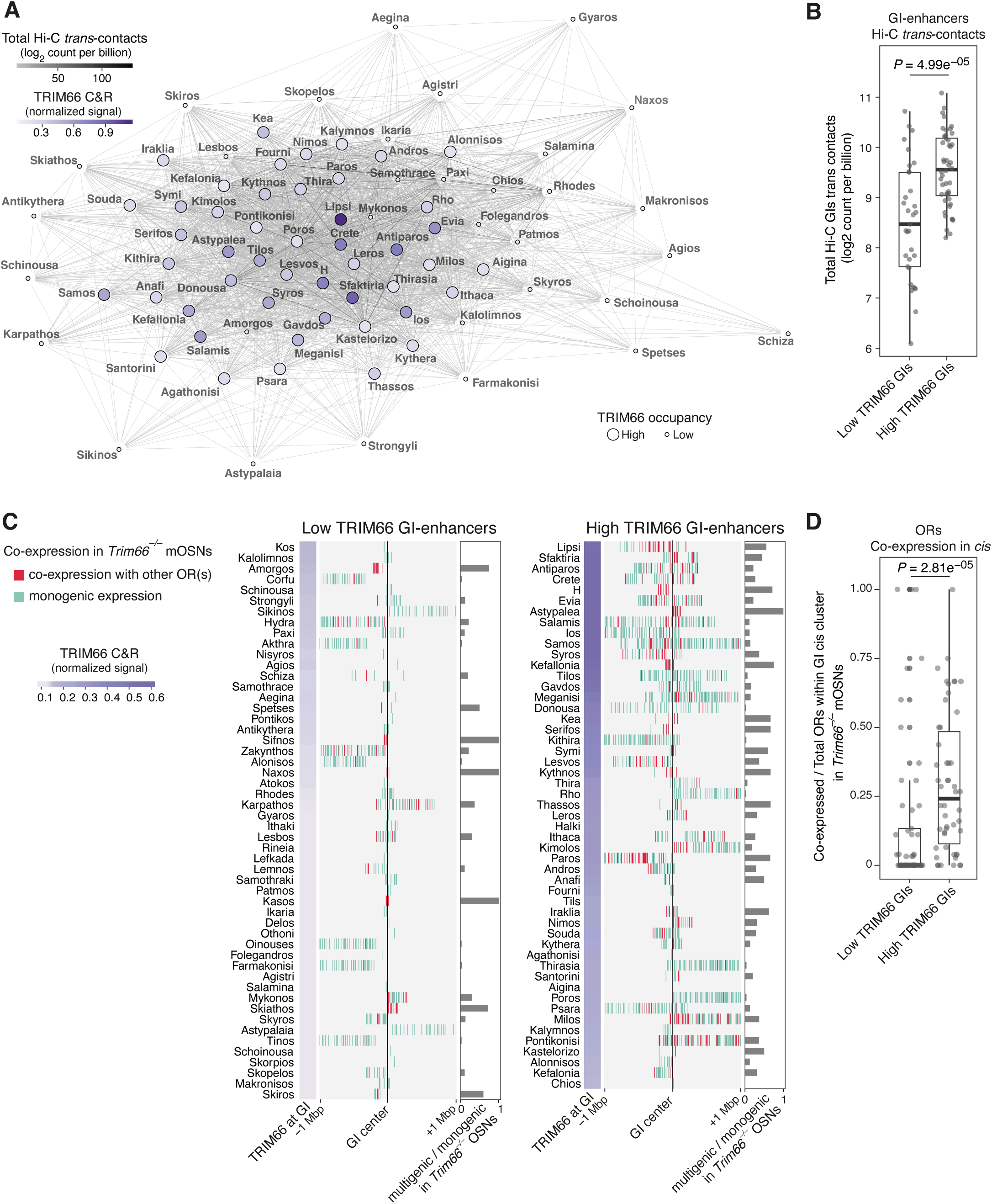
TRIM66 binds to *trans*-interacting GI-enhancers and represses OR genes in *cis*. (A) Network representation of GI-enhancers trans-contacts integrating Hi-C data (GSE243479, GSE243480) and TRIM66 occupancy determined by CUT&RUN. (B) Total Hi-C trans contacts of TRIM66-occupied GI-enhancers (n=48) *versus* GI-enhancers with low TRIM66 occupancy (n=32). TRIM66-bound GI-enhancers displayed a higher propensity to make trans interactions than those devoid of TRIM66 (*P*=0.499e-5, Wilcoxon-Mann-Withney U-test). Each dot represents a GI, horizontal lines show median values, boxes represent the 25^th^ to 75^th^ interquartile range, whiskers extend from the 75^th^ quartile + 1.5 * interquartile range (that is, the distance between the 75^th^ and 25^th^ quartiles) and the 25^th^ quartile - 1.5 * interquartile range. (C) Heat map of OR genomic clusters centered on GI-enhancers and sorted by TRIM66 occupancy. Left panel: TRIM66 low-occupancy, right panel: TRIM66 high-occupancy. Clusters were tiled in 10 kb non-overlapping windows. Windows containing multiple OR genes were counted once. OR expressed with other ORs in *Trim66*-mutant mOSN are shown in red. (D) Ratio of coexpressed OR in *cis* of GI-enhancers devoid of TRIM66 versus ratio of coexpressed OR in *cis* of TRIM66-occupied GI-enhancers. The ratio of coexpressed OR genes in *Trim66*-deficient mOSN over the total number of OR genes within the corresponding cluster was calculated. OR located in *cis* of a TRIM66-bound GI-enhancers are more likely to become expressed with other GI-enhancers after *Trim66*’s removal (*P*=2.81e-05, Wilcoxon-Mann-Withney U-test). Each dot represents a GI, horizontal lines show median values, boxes represent the 25^th^ to 75^th^ interquartile range, whiskers extend from 75^th^ quartile + 1.5 * interquartile range (that is, the distance between the 75^th^ and 25^th^ quartiles) and the 25^th^ quartile - 1.5 * interquartile range.

### Disorganized OSNs axonal projections in *Trim66*-deficient mice

Does the abnormal expression of multiple ORs per neuron influence their connectivity with the OB? The subpopulation of OSNs expressing the same OR normally project their axons to one of two possible glomeruli in the OB ^20^. OR gene swap experiments have demonstrated that the identity of the expressed OR instructs, via an unknown mechanism, the location of the glomerulus to which the OSN projects ^8^. To our knowledge, *Trim66* loss-of-function mutations is the only known case of widespread multigenic expression of OR genes. We used this model to test the requirement of singular OR expression for proper axonal guidance. We visualized the axons of OSN expressing *Olfr446* using our *Olfr446^T2i-TAU-GFP^*reporter allele. As expected, OSNs expressing *Olfr446* project their axons to two specific glomeruli of the OB (Figure 4A, left panel). In contrast, *Trim66* loss-of-function causes a massive disorganization of axonal projections of *Olr446*-expressing mOSNs that were unfocused and invaded large areas of the OB (Figure 4A, right panel). The data show that the expression of a single OR per OSN is necessary for proper axon guidance.

**Figure 4:**
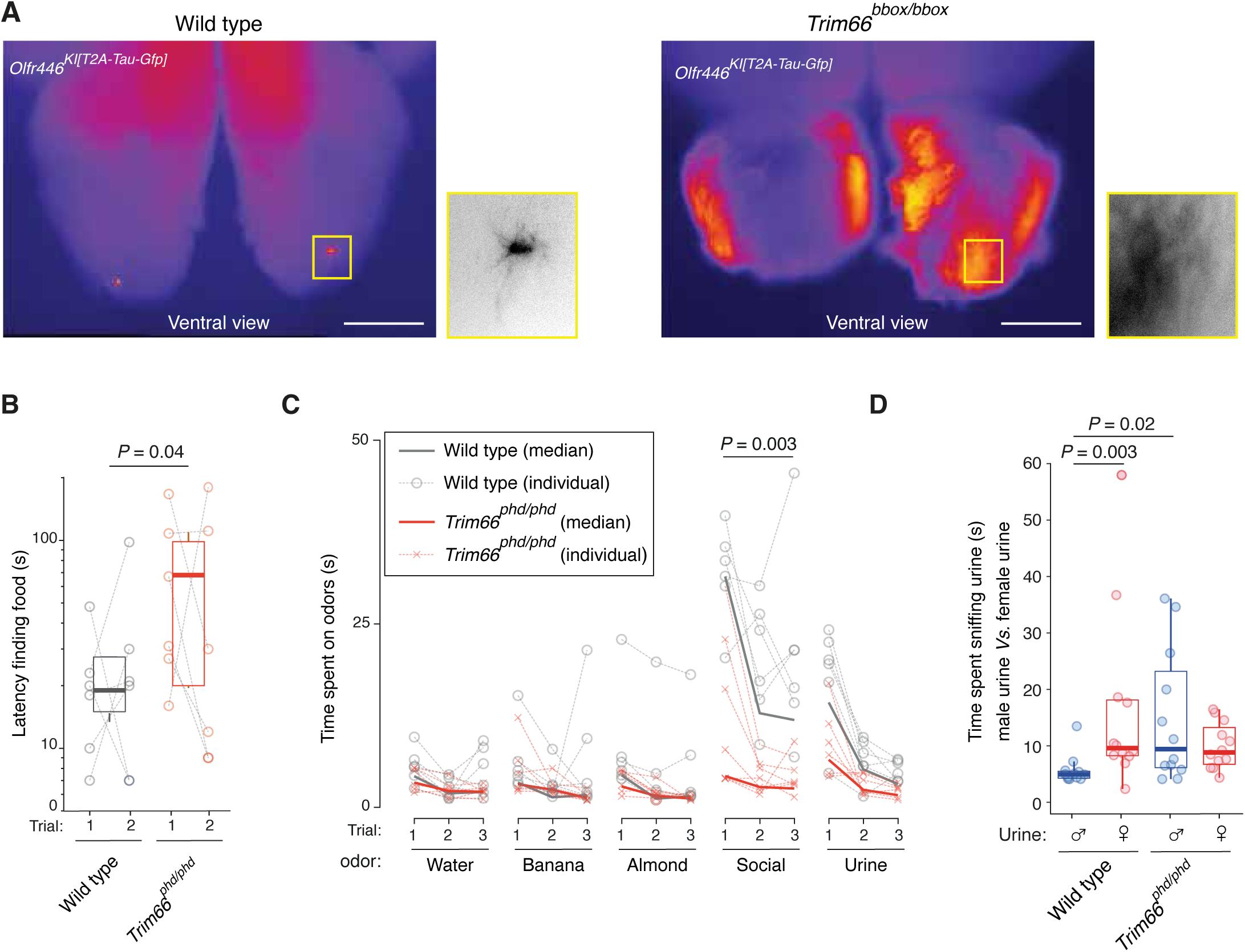
Disruption of the olfactory map in the bulb and impairment of odors’ discrimination. (A) Representative whole-mount images of the olfactory bulb (ventral view) with staining of *Olfr446*-expressing mOSN axons. The axons converge into a single glomerulus in the WT olfactory bulb (left). In contrast, *Olfr446*-expressing mOSNs disrupted for *Trim66* project their axons in a disorganized and unfocused manner to numerous glomeruli. Signal is displayed using fire LUT to facilitate the visualization of signal intensity, where blue is the minimum and white is the maximum. Central panels: zooms of WT and mutant glomeruli shown in the yellow boxes. Scale, 1000 µm. (B) Buried food olfactory test. *Trim66^phd/phd^* (n=6) displayed a significantly higher latency to find a piece of food hidden under the bedding than for WT animals (n=6; *P*=0.04, Mann–Whitney U test). Each data point represents a test animal, with a gray line connecting the data from the same animal. The box plot shows the first, second, and third quartiles, with whiskers indicating the non-outlier data range. (C) Olfactory habituation/dishabituation test. *Trim66^phd/phd^* male mice spent significantly less time sniffing social odors (cage smell) than WT animals (*n*=6 for each genotype; Mann–Whitney U test). No significant differences were observed in the time spent for other odors tested, including neutral (water) or female urine odors. The odors were presented in series, each presented three times. (D) Sex urinary odor preference test. *Trim66^phd/phd^* and WT males (n=6) were presented with male and female urines, and the time spent sniffing each urine sample was recorded. *Trim66^phd/phd^*males displayed no preference for female over male urine and investigated male urine significantly longer than WT males did. The box plot shows the first, second, and third quartiles, with whiskers indicating the non-outlier data range, and *P*-values < 0.05 are shown (Mann–Whitney U test).

### *Trim66* loss-of-function impairs olfaction

How does the impairment of the OSN wiring pattern affect olfactory-related behavior? To address this question, we performed three different behavioral assays on adult male WT and *Trim66*-deficient mice. First, we examined the animal’s ability to locate food hidden beneath bedding (buried food test)^21^. The test was conducted twice on each animal across different days. In both trials, *Trim66^phd/phd^* mice showed significantly prolonged latencies to find the food compared to WT mice (Figure 4B). We next performed an olfactory habituation/dishabituation test to examine responses to a wider variety of odorants ^22,23^. WT animals generally showed robust responses, such as sniffing, when a novel odor was presented, but the response became weaker when the same odor was repeatedly presented (Figure 4C). In particular, WT mice showed stronger responses to social odor (cage scent or urine) than to non-social odor (isopentyl acetate or benzaldehyde) or pure water (negative control). In contrast, *Trim66^phd/phd^* mice did not show such preference for any particular smell, leading to significantly less time spent near the smell than WT mice did. To further investigate deficits in social olfactory behavior due to TRIM66 deficiency, we performed a two-choice odor-preference test using male and female urine samples (Figure 4D). As previously reported ^22^, WT males were more attracted to female urine samples than to male ones. However, *Trim66^phd/phd^* males showed no preference and spent more time sniffing the male urine sample than WT males. Taken together, the outcome of these behavior tests revealed impaired olfaction in *Trim66* loss-of-function mutations, likely due to co-expression of multiple OR genes and the resulting mistargeting of mOSN axons to the olfactory bulb.

## DISCUSSION

We identified TRIM66 as a chromatin protein necessary for preventing the expression of more than one OR gene in olfactory neurons. The data presented here demonstrate that the exclusive transcription of a single OR gene per neuron is enforced by epigenetic silencing rather than by the selective activation of OR genes. We suggest a mechanism that depends on the recruitment of TRIM66 to a subset of GI-enhancers with *cis*-activation potential. We speculate that TRIM66 could be the chromatin feature that distinguishes transcriptionally silent GI-enhancer aggregates (inactive GI-hubs) from the unique transcriptionally active GI-enhancer hub within a cell ^12,19^. The molecular cues that target TRIM66 to GI-enhancers remain unknown. TRIM66 lacks any known DNA binding domain, yet it could be targeted to GI-enhancers via its binding to an unknown transcription factor, similarly to its paralogue TRIM28, also known as KAP1, which is recruited to parasitic DNA sequences via its interaction with transcription factors belonging to the large family of Krüppel-associated box (KRAB)-containing zinc finger proteins ^24^. It is noteworthy that the function of these two paralogue chromatin proteins—TRIM28 and TRIM66—lies in the functional partitioning of the mammalian genome by silencing selected promoters: retrotransposons for TRIM28 and OR genes for TRIM66 ^25^. Hence, these two epigenetic silencing mechanisms may have evolved from an ancestral TRIM-dependent repressive mechanism.

## METHODS

### Data and code availability

All data are available in the main text and in the supplementary materials, and all reagents can be made available from the corresponding author upon request.

All code is available in the GitHub repository: https://github.com/boulardlab/trim66-moe The publicly available datasets reanalyzed are described in Table S4.

Sequencing data have been deposited in ArrayExpress under the following accessions:

- Bulk RNA-seq: E-MTAB-15645
- Single-cell 10X RNA-seq whole MOE: E-MTAB-15996
- CUT&RUN: E-MTAB-15644
- Single-cell Smart-seq2 RNA-seq purified *Olfr446*-expressing cells: E-MTAB-16019

### Animal care and handling

Mouse husbandry and experiments adhered to European and national regulations for the protection of vertebrate animals used for experimental and other scientific purposes (directives 86/609 and 2010/63). All protocols were approved by the EMBL Animal Care and Use Committee under protocols 22-016_RM_MB and 24-011_RM_MB and the Italian Ministry of Health License under protocol n. 473/2025-PR. Mice were housed in the pathogen-free Animal Care Facility at EMBL Rome on a 12-hours light–dark cycle in temperature- and humidity-controlled conditions with free access to food and water.

### Main olfactory epithelium collection

Experimental animals were euthanized by cervical dislocation, taking care not to aspirate the blood into nasal passages by damaging the veins and arteries in the neck. The skull was collected, and using surgical tools under a dissecting microscope, the skull bones surrounding the nasal tissue were removed. The entire and intact main olfactory epithelium was dissected and transferred into ice-cold PBS and then processed for the downstream experiment.

### Generation of murine alleles: Trim66^phd^, Trim66^bbox^, Trim66^HA^, Olfr446^T2A-TAU-GFP^

The *Trim66^phd^* allele was previously produced in the Boulard laboratory ^17^. The other alleles were created by CRISPR/Cas9-editing using C57BL/6J (Charles River) zygotes following a previously described strategy ^26^. The *Trim66^bbox^* allele consists of an insertion of a premature stop codon in exon 4 (*Trim66-202*) that encodes a Bbox domain; the sequence of the CRISPR crRNA oligo was TCTGCACATACTGCAACCGC. The premature stop codon was inserted in exon 3 transcript *Trim66-201*, (ENSEMBL v91); genomic coordinate: Chr 7. 109083675 (GRCm39/mm39). For *Trim66^HA^*, the crRNA oligo sequence CTGACTTACCCTGGCCATAG was used to insert a 3xHA tag at the N-terminus of transcript *Trim66-201*; genomic coordinate: Chr 7. 109085259. For *Olfr446^T2A-TAUGFP^*, the crRNA oligo sequence GTGCCTTGAGACGAGTTCTT was used to insert a tau-GFP tag with a ribosomal skipping 2A linkage peptide (termed T2A) at the C-terminus of transcript *Or2a12-201*; genomic coordinate: Chr 6. 42905070. For *Trim66^bbox^* and *Trim66^HA^*, annealed sgRNAs were complexed with Cas9 protein and combined with their respective lssDNA donors (Cas9 protein 20 ng/mL, sgRNA 20 ng/mL, lssDNA 10 ng/mL, all IDT). These reagents were microinjected into zygote pronuclei using standard protocols ^27^. For *Olfr446^T2A-TAU-GFP^*, zygotes were infected with rAAVs containing the required knock-in sequences and then electroporated with the sgRNA/Cas9 RNP complex ^26^. In all cases, after overnight culture, the 2-cell stage embryos were surgically implanted into the oviduct of day 0.5 post-coitum pseudopregnant CD1 mice.

Founder mice were screened for by PCR, first using primers flanking the sgRNA cut sites, which identifies InDels generated by NHEJ repair and can also detect larger products implying HDR. Secondary 5’ and 3’ prime PCRs using the same primers in combination with template-specific primers allowed for the identification of potential founders; these PCR products were then Sanger sequenced and aligned with the in silico design. Sequences of the single-stranded donor templates are provided in Table S2. The strains were maintained on a C57BL6/J genetic background. Transgenic mouse production was performed by the Gene Editing & Virus Facility at EMBL.

### Olfactory behavioral assays

Adult males homozygous for *Trim66^phd^* (n=6), *Trim66^bbox^*(n=6) and littermate control animals (n=6+6) were subjected to three olfactory behavior tests: 1) a buried food test, where the time to find a piece of food buried under bedding was measured ^21,28^; 2) a habituation/dishabituation test, where the time spent sniffing novel odors was measured ^21–23^; and 3) a two-choice odor preference test, where the time spent sniffing male or female urine samples was measured and compared ^22^. For each assay, the test animal was transferred to a clean cage (length, 46 cm; width, 23.5 cm; height, 20 cm) and adapted to the new environment for 30 min/day for 5 days before the experiment. All experiments were video recorded (720-by-1280 pixels; 24 or 30 Hz) and manually analyzed in a double-blind manner.

For the buried food test, the subject animals were food-deprived the day before the test. A piece of food was buried approximately 1 cm below the bedding at a random corner of a clean cage. The subject mouse was then placed in the middle of the cage, and the time spent by the animal to find the buried food by smell was measured.

For the habituation/dishabituation test, the animal was exposed to five different odors (40 μL soaked in a cotton swab), each presented three times consecutively, in the following sequence: water, isopentyl acetate (“banana” odor; Sigma, 306967), benzaldehyde (“almond” odor; Sigma, W212709), social cage odor (obtained by scratching a cotton swab on cage floors housing multiple mice), and female urine (collected from multiple animals). In each trial, the odor-containing cotton swab was inserted from the top of the cage (about 6 cm from the short side of the cage, 12 cm from the long side, and 8 cm from the floor). The time spent sniffing the swab in each trial (duration, 3 min) was measured.

For the two-choice odor preference test, 40 µL of urine collected from multiple animals of the same sex was soaked into cotton swabs. The male and female urine swabs were fixed to a cage lid approximately 10 cm apart. The cage lid was then placed over a clean cage containing a male test animal. The time spent sniffing female and male urine was measured, respectively (5 min/trial).

### Western blot

Immunodetection of TRIM66 was performed on nuclear extracts isolated from MOEs dissected from 10–15 weeks-old mice of the following genotypes: WT, *Trim66^phd/phd^*,*Trim66^bbox/bbox^*, *Trim66^phd/bbox^*. The nuclei isolation was performed as previously described ^29^. In brief, the MOE was rapidly dissected in ice-cold homogenization buffer (0.25 M sucrose, 25 mM KCl, 5 mM MgCl₂, 20 mM Tricine-KOH, 1 mM DTT, 0.15 mM spermine, 0.5 mM spermidine). The tissue was minced and homogenized with a loose pestle in 5 mL of homogenization buffer. Subsequently, 300 μL of 5% IGEPAL-630 solution was added, and the homogenate was further processed with five strokes of a tight pestle. The homogenate was filtered through a 40 µm cell strainer, layered over a gradient composed of 50%, 40%, and 30% OptiPrep™ Density Gradient Medium (Sigma-Aldrich, D1556), and centrifuged at 10,000 g for 18 minutes at 4 °C in a swinging bucket rotor. Nuclei were collected from the 30%-40% interface and incubated in RIPA buffer (150 mM NaCl, 1% NP-40, 0.5% sodium deoxycholate, 0.1% SDS, 50 mM Tris-HCl, pH 8.0) until fully solubilized. The protein lysates were diluted to the appropriate concentration in SDS loading buffer (200 mM Tris–HCl, pH 6.8, 8% SDS, 40% glycerol, 400 mM DTT, 0.4% bromophenol blue; all from Sigma-Aldrich) and denatured at 95 °C for 5 minutes. Samples were separated using 4–12% Bis-Tris gels (Invitrogen) in MOPS buffer and transferred onto PVDF membranes using the Trans-Blot Turbo Transfer System (Bio-Rad) for 13 minutes. Membranes were blocked in 5% non-fat dry milk (Roth) in PBS containing 0.1% Tween-20 (Sigma-Aldrich) for 1 hour at room temperature. Primary antibodies were incubated overnight at 4 °C in blocking buffer. The anti-TRIM66 antibody (EMBL PEPcore ^17^) was used at 1:50, and Lamin A/C (sc-376248; Santa Cruz Biotechnology) at 1:2000. Membranes were then washed five times for 5 minutes each in washing buffer I (0.5% Triton X-100, 0.5 M NaCl, 1X PBS), followed by a 10-minute wash in washing buffer II (0.5 M NaCl, 1X PBS), and a final 15-minute wash in PBS. HRP-conjugated secondary antibodies were incubated for 1 hour at room temperature in blocking buffer. Membranes were washed five times for 5 minutes each in TBS-T, and signal detection was carried out using ECL substrate (GE Healthcare) according to the manufacturer’s instructions. Images were acquired using the Amersham ImageQuant 800 (GE Healthcare). The regions corresponding to relevant molecular weights were cropped for figure presentation.

### Histological sections and immunofluorescence microscopy

The MOE was isolated from adult mice, fixed in 4% paraformaldehyde, and embedded in paraffin. The 8 μm coronal sections were dewaxed in xylene twice for 10 min and then rehydrated in the ethanol series of two washes in 100% EtOH for 5 min and a single wash in 95% EtOH for 2 min, 70% EtOH for 2 min, and 50% EtOH for 2 min, followed by two washes in distilled water for 5 min before heat/antigen retrieval step. The antigen retrieval was performed in the microwave (730 W) for 20 min in 10 mM Tris-EDTA buffer pH 9. Cooled slides were washed in PBS twice for 5 min, then permeabilized with 0.3% Triton X-100 in TBS for 10 min. The slides were washed three times for 5 min in 0.1% TritonX-100 in TBS. The blocking of the slides was done in 5% natural donkey serum in the TBS buffer with 0.1% Triton X-100 for 30 min at RT. The slides with antibodies were incubated overnight at 4 °C in the blocking solution. The HA antibody (Roche, #11867423001) was used at a dilution of 1:100. The following day, the detection (for the IF experiment) was performed using an avidin/biotin-based peroxidase system (VECTASTAIN Elite ABC system, anti-Rat, #PK-6104), with a secondary streptavidin Alexa Fluor 546-conjugated (Invitrogen). Then, the cell-marker antibodies were incubated overnight at 4 °C in the blocking solution. The HA detection by IHC was performed similarly using peroxidase deposition of diaminobenzidine (DAB) precipitate. The antibodies anti-GAP43 (AB5220, Sigma-Aldrich) and anti-OMP (ab183947, Abcam) were diluted at 1:100 in the blocking solution. GAP43 (iOSNs) and OMP (mOSNs) were detected with a secondary goat anti-rabbit antibody, Alexa Fluor 647 (Thermo Fisher Scientific) at 1:500 dilution. The slides were mounted with Fluoromount G (Thermo Fisher Scientific). The imaging was obtained with confocal microscopy at 60X magnification with a Nikon AX microscope with Galvano scanner. Image processing was carried out using ImageJ, with adjustments made to brightness, contrast, and levels of separate channels for composite images.

### Imaging the whole-mount olfactory bulb

Adult animals (12 weeks-old) were anesthetized deeply with a mix of ketamine and xylazine (100 mg/kg + 10 mg/kg) and perfused transcardially with 4% PFA/PBS. The following genotypes were analyzed: *Trim66^WT^*; *Olfr446^KI[T2A-TAU-GFP]^*(n=3), *Trim66^phd/phd^ Olfr446^KI[T2A-TAU-GFP]^* (n=1) ; *Trim66^bbox/bbox^* ; *Olfr446^KI[T2A-TAU-GFP]^* (n=2). Brains were dissected and postfixed in 4% PFA/PBS overnight at 4 °C. The following day, olfactory bulbs were dissected and imaged using a Leica Thunder DMi8 inverted microscope, N Plan 5x NA 0,12 objective. LED 475 nm 40%, exposure time 500 ms, gain 6.01. Tile imaging was acquired using LAS X Navigator with a 20% overlap.

### RNA-FISH

RNA was detected by Hybridization Chain Reaction (HCR) combined with fluorescence *in situ* hybridization (RNA-FISH). Five week-old mice were sacrificed by cervical dislocation. The olfactory epithelia were dissected, embedded in Tissue-Tek O.C.T. compound (Sakura; #4583), and rapidly frozen on dry ice. Frozen tissue blocks were sectioned using a cryostat (Leica CM 3050S). The sections of 5 µm thickness were placed on glass slides and stored at –80 °C until further use. RNA-FISH was performed using *in situ* hybridization chain reaction HCR™ RNA-FISH (v3.0) technology from Molecular Instruments according to the manufacturer protocol for fresh/fixed frozen tissues. The sets of HCR probes for the two OR transcripts: *Mouse_Olfr446* (Gene ID: 258292) and *Mouse_Olfr231* (Gene ID: 404222) were generated by Molecular Instruments. Images were acquired with a V3 X-light spinning disk (Crest Optics) confocal, which also has epifluorescence and DeepSIM super-resolution modes available, built around a Nikon Ti2 microscope body. It has a 7-line (405, 446, 488, 518, 545, 637 and 748 nm) laser engine Celesta (Lumencor) available, and a Prime 95B sCMOS camera (Photometrics). The whole system is managed by NIS-Elements (Nikon). The system is equipped with the following objectives: a 4x, a 10x, a 20x NA 0.75, a 40x dry NA 0.95 and a 60x NA 1.42 oil immersion objective. Whole-tissue images were acquired with the 20x, while multiple regions of interest were taken with the 60x objective.

### Bulk RNA sequencing assay

RNA was extracted using TRIzol (Invitrogen) from entire MOEs dissected from 10-15 week-old WT (n=6) and *Trim66^phd/phd^* animals (n=6). For each sample, 1 µg of total RNA was then treated with DNase (Invitrogen Ambion TURBO DNase) at 37 °C for 15 minutes. Subsequently, RNA was purified by precipitation using 10% 3 M sodium acetate (NaAc), 1 μl of glycogen, and 2 volumes of absolute ethanol (EtOH), followed by a wash with 75% EtOH. The integrity of the RNA was assessed using the Tape Station system (Tape Station RNA HS D1000 Kit). Stranded mRNA-seq library preparation was performed by the EMBL Genomics Core Facility using the NEBNext® Ultra™ II RNA Library Prep Kit for Illumina using the NEBNext Poly(A) mRNA Magnetic Isolation Module. The mRNA-seq libraries were sequenced on the Illumina NextSeq 500 platform using the paired-ends sequencing mode.

### 10X Single-cell RNA sequencing assay

MOEs were isolated from WT (n=2), *Trim66^phd/phd^* (n=1), *Trim66^bbox/bbox^* (n=1) adult animals (18-19 weeks old). The tissue was minced and dissociated into single-cell suspension following the instructions from the Papain Dissociation System (Worthington) with an enzymatic dissection time of 30 min at 37 °C. After the last step, cells were resuspended in sorting buffer (1% PBS with 5% FBS, supplemented with 1X glucose solution and 10X sodium pyruvat), then stained with 2 µg/mL propidium iodide for 10 min at RT. The prepared sample was filtered through a 50 µm nylon filter, then sorted using a BD FACS Aria II Flow Cytometer (BD Biosciences) for the target of 150,000-200,000 events. Sorted living cells were centrifuged to enrich the cell concentration (5 min, 300 x g), then resuspended in a smaller volume. The cells were quantified using the Moxi cell counter (Orflo, S cassettes), then brought to a target concentration of ∼1000 cells/µl. The single-cell sorting, labeling, and library preparation were performed using the 10X Genomics instrument and reagents (Chromium Controller X with Chromium Next GEM Single cell 3’ Reagent kit v3.1 Dual index, PN-1000269, PN-1000215, and PN-1000127) according to the manufacturer’s instructions (version CG000315 Rev C). For the *Trim66^phd/phd^* and corresponding WT samples, a target cell number of 8,000 was aimed for; for the *Trim66^bbox/bbox^*samples a target concentration of 10,000 cells. Quality control and concentration measurements of the samples during library preparation were performed with the High Sensitivity dsDNA kit (Qubit) and the D5000 High Sensitivity TapeStation reagents (Agilent). The mRNA-seq libraries were sequenced by the EMBL Genomics Core Facility on the Illumina NextSeq 2000 platform using the P3 100 bp paired-end sequencing mode (1,100 MIO clusters).

### Smart-Seq2 single cell RNA sequencing assay

MOEs from two *Olfr446^T2A-TAU-GFP/+^; Trim66^wt^* and two *Olfr446^T2A-TAU-GFP/+^; Trim66^bbox/bbox^* animals (5 weeks old) were isolated as described before. The tissue was minced and dissociated into single-cell suspension following the instructions from the Papain Dissociation System (Worthington) with enzymatic digestion time of 45 min at 37 °C ^30^. After the last step, cells were resuspended in sorting buffer (1% PBS with 5% FBS, supplemented with 1X glucose solution and 10X sodium pyruvate). The prepared samples were filtered through a 50 µm nylon filter, then sorted using a BD FACS Aria II Flow Cytometer (BD Biosciences). OSNs were sorted directly into a TrimTech 96 well plate (Eppendorf # 951020401) containing lysis buffer (Triton X100 (0.5% (v/v), Sigma (#T8787), 1 µl RNAse inhibitor (40 U), Clontech Takara (#2313A), 1 µl dNTPs (10 mM, Kappa (#KK1017) and 1 µl tailed oligo-dT30VN 5′AAGCAGTGGTATCAACGCAGAGTACT30VN-3′ (5 µM Eurofins); total volume 4.4 µl). Upon sorting, OSNs were snap frozen and stored at −80 °C until further processing. Single-cell full-length cDNA libraries for mRNA sequencing were prepared by the EMBL Genomics Core Facility using a modified Smart-Seq2 protocol described in ^18^. Reverse transcription was performed using the SuperScript IV Reverse Transcriptase kit (ThermoFisher #18090200) with the following reaction: 2 μL SSRT IV 5x buffer, 0.5 μL 100 mM DTT, 2 μL 5 M betaine, 0.1 μL 1 M MgCl_2_, 0.25 μL 40 U/μL RNAse inhibitor (Takara #2313A), 0.25 μL SSRT IV, 0.1 μL 100 µM TSO, 1.15 μL RNase-free H_2_O; thermal conditions: 52 °C 15’, 80 °C 10’. The pre-amplification reaction was performed using the KAPA HiFi HotStart ReadyMix (Roche, #KK3604) with a number of 16 PCR cycles (98 °C for 20 sec, 67 °C for 20 sec, 72 °C for 6 min, 4 °C hold). Amplified DNA was cleaned-up using 0.6x SPRI beads (Beckman, #B23318). Tagmentation procedure was performed with the Tn5 Transposase purified by the EMBL Protein Expression and Purification Core Facility as previously described by ^31^. The reaction was performed on a 100 pg normalized input sample with the following conditions: 55 °C for 8 min, 4 °C hold. Next, the enrichment PCR reaction was carried out with the following conditions: 12-21 PCR cycles, 98 °C for 20 sec, 58 °C for 15 sec, 72 °C for 30 sec, 72 °C for 3 min, 10 °C hold. The final pool, consisting of 5-8 µl of each sample without normalization, was further cleaned up with 0.8x SPRI beads (Beckman, #B23318). Quality control of the library was performed using an Agilent 2100 Bioanalyzer and High Sensitivity DNA kit (Agilent Technologies, # 5067-5579). The sample pool was sequenced by the EMBL Genomics Core Facility in one run using the Cloudbreak kit (medium output 2×150) on an AVITI sequencer.

### CUT&RUN assay

The CUT&RUN profiling experiment was performed in biological triplicate. The MOE from WT and *Trim66^HA/HA^* animals of 10-15 weeks-old were dissected, and the nuclei isolation was performed as previously described ^32^. For each condition, 1 million nuclei were processed following the live nuclei CUT&RUN protocol ^33^. The nuclei were incubated for 10 min at RT under gentle rotation with 25 µL of Concanavalin A-coated magnetic beads (Bangs Laboratories #BP531) previously washed twice in Binding Buffer (20 mM HEPES–KOH pH 7.9, 10 mM KCl, 1 mM CaCl_2_, 1 mM MnCl_2_). The bead-bound nuclei were isolated on a magnetic stand and blocked with 1 mL of Blocking Buffer (20 mM HEPES pH 7.5, 150 mM NaCl, 0.5 mM Spermidine, 0.1% BSA, 2 mM EDTA, 1X cOmplete Mini EDTA-Free Protease Inhibitors) for 5 min at RT. After blocking, the beads were washed in Wash Buffer (same as Blocking Buffer without EDTA) and resuspended in 500 μL of Antibody Buffer containing 1 μg of anti-HA antibody (Abcam, ab9110). Samples were incubated overnight at 4 °C with gentle rotation. The next day, samples were washed with Wash Buffer and resuspended in 300 μL of Wash Buffer supplemented with 700 ng/mL pA/G MNase. After 1 hour of incubation at 4 °C under gentle rotation, samples were washed twice and resuspended in 50 μL of Wash Buffer, then placed on ice to pre-cool to 0 °C. The pA/G MNase protein was produced by cloning a synthetic gBlock (IDT) into the pETM14 vector, followed by bacterial expression and purification at EMBL PepCore. Targeted chromatin digestion was initiated by adding 2 μL of 100 mM CaCl_2_ and was incubated for 30 min at 0 °C. The digestion reaction was stopped by adding 50 μL of 2X Stop Buffer (200 mM NaCl, 20 mM EDTA, 4 mM EGTA, 1% IGEPAL, 1 mM MnCl_2_). Normalization Drosophila Spike-In was added to the samples in Stop Buffer. Samples were incubated at 37 °C for 15 min to release the CUT&RUN fragments from the insoluble nuclear chromatin and centrifuged at 16,000 g for 5 min at 4 °C. Supernatants were collected after magnetic separation and transferred to new tubes. Samples were incubated at 70 °C for 10 min supplemented with 2 μL of 10% SDS and 2.5 μL of 20 mg/mL of Protein K. DNA fragments were then purified and size-selected using SPRIselect magnetic beads (Beckman Coulter #B23318) according to the manufacturer’s protocol for single selection to purify fragments > 100 bp. DNA fragments were eluted in 30 μL of nuclease-free water. DNA concentrations were quantified using Qubit and libraries were prepared with the NEBNext Ultra II DNA Library Prep Kit for Illumina (E7645S) following the manufacturer’s protocol. Libraries were size-selected from Nusieve 3:1 Agarose gel (Lonza) to isolate fragments between 200-500 bp, followed by purification using the Monarch DNA Gel Extraction Kit Protocol (New England Laboratories #T1020). The quality of the libraries was assessed using the Tape Station system (Tape Station DNA HS D1000 Kit). Sequencing was performed on the Illumina NextSeq 500 platform.

### Bulk RNA-seq analysis

Bulk RNA-seq data were analyzed with the 3T-seq Snakemake pipeline ^34^. In short, the pipeline performs adapter trimming, reference genome mapping, duplicated reads removal, gene expression quantification, and differential expression analysis of both protein coding genes, transposable elements, and tRNAs. Adapter trimming is performed with Trimmomatic v0.39, reference genome mapping and gene expression quantification with STAR v2.7.6a, duplicates removal with Picard v2.27.4 and differential expression testing with DESeq2 v1.30.1. The Gencode M25 build of the GRCm38 (mm10) mouse genome sequence was used. The 2021 version of the Mouse Genomics Informatics (MGI) genome annotation was used. For transposable elements, the mm10 RepeatMasker track was downloaded from the UCSC web server using the built-in functionality of the 3T-seq pipeline. For differential expression testing, the Wald t-test was used. Adjusted *P*-value cutoff was set to 0.01 and a log2 fold change threshold of 0.5 was set.

### 10X scRNA-seq analysis

For each sequencing library, a barcode-features matrix of gene counts was obtained using Cell Ranger version 6.1.1. Mouse reference files (version 2020-A) were downloaded from the 10X Genomics website. Downstream analysis was performed with Seurat version 4.1.0 running on R version 4.1.0.

Each pre-filtered barcode-feature matrix was imported into the R environment. Percent of mitochondrial RNA contamination was computed from genes whose name starts with “mt-”. To remove empty droplets and doublets, barcodes with less than 200 or more than 5000 detected features were discarded. In addition, cells showing more than 10% mitochondrial RNA content were discarded, as high mitochondrial RNA concentration is associated with cellular damage. This filtering procedure resulted in 3016 *Trim66^phd/phd^*, 7214 *Trim66^bbox/bbox^* and 2395 and 4020 WT cells for *Trim66^phd/phd^* and *Trim66^bbox/bbox^* -paired samples respectively.

For each remaining cell, a score associated with cell cycle was calculated. Human gene lists specific to G2M (54 genes) or S (43 genes) phase were retrieved from Seurat data files. Their human identifiers were converted to mouse gene symbols using the biomaRt package version 2.62 from Bioconductor project ^35^. Fifty one out of fifty four and 40/43 identifiers were successfully mapped for G2M phase and S phase respectively. A score was computed for each phase using the CellCycleScoring function from Seurat. Their difference was calculated and used downstream to model cell cycle effect.

The sctransform method was used to normalise each count matrix separately ^36^. This is a regularised regression method based on the negative binomial model. It corrects both for technical and biological sources of variation. For each gene, the method first fits a general linear model to remove the effect of library size, then runs a kernel density estimate to regularise the model parameters and finally a second round of regression with parameters constrained to the values learned earlier is performed. Gene residuals are interpreted as normalised expression values. This framework is flexible enough to accept additional covariates. Mitochondrial RNA percentual content and the cell cycle score previously computed were used as biological covariates and their effect modelled and removed during the normalisation procedure.

*Trim66^phd/phd^*, *Trim66^bbox/bbox^* and wild type samples were integrated using Canonical Correlation Analysis. Top 3000 variable genes were detected across the four samples and used to find the anchors needed for integration. The integrated dataset is composed of 3000 genes and 16654 cells. Principal component analysis was performed, the first 21 principal components were calculated and used to compute the uniform manifold approximation and projection (UMAP). Clusters of cells were detected using the shared nearest neighbour with modularity optimization algorithm set to Louvain algorithm with multilevel refinement as implemented in the Seurat package; clustering resolution was set to 1.5. The procedure resulted in 36 clusters. Cluster marker genes were detected with the FindAllMarkers function from the Seurat package. For main olfactory epithelium cell populations, marker genes were manually curated from literature (Table S1). By assessing expression of manually curated markers and automatically detected ones, cluster labels were manually inferred.

For each immediate neural precursor (INP), immature olfactory sensory neuron (iOSN) and mature olfactory sensory neuron (mOSN) individual cell, the number of expressed OR genes was computed as follows. A receptor was considered expressed if displaying a normalized read count (Seurat slot “counts”, assay “SCT”) greater than 5. For each cell, the total number of expressed receptors was calculated by summing the number of OR genes passing the filter above. This information was used to calculate histograms (Figure S3B,C) and to color cells in the UMAP representation (Figure 1D,E and S3A).

### Smart-Seq2 scRNA-seq analysis

Raw sequencing files were aligned to the mouse reference genome (GRCm38) using the 3t-seq pipeline with default parameters ^34^. Tabular output files were loaded into R (v. 4.4.1) and combined into a single per-cell per-gene count matrix, selecting the unstranded gene counts.

Downstream analysis was performed with Seurat (v. 5.3.0) ^37^. Raw gene counts were stored as “RNA” layer of the Seurat object, and genes with zero-counts were removed from downstream analyses. Mitochondrial and ribosomal count percentage was evaluated per single cell, using genes starting with “mt-” and “Rpl”, respectively. Cells with a mitochondrial percentage greater than 10 were removed as stressed cells; cells with less than 300 features were also discarded. This resulted in 25461 features across 459 cells. Additionally, 27 cells, not expressing *Olfr446* were removed from further analyses. SCTransform was employed to normalise the data, with default parameters and without regressing out any covariate ^36^. The top 3,000 variable features were identified based on average expression across cells and standardised variance (VariableFeaturesPlot, Seurat). To perform dimensionality reduction of the dataset, a PCA analysis was performed, and subsequent UMAP coordinates were calculated using the top 14 dimensions. Cell cycle scores were evaluated using a set of canonical marker genes for the S and G2M phases, provided by Seurat. These were converted to the mouse orthologs using the g:Orth tool.

Cell type annotation was performed based on a manually created gene list with markers for the cell types present in the mouse olfactory epithelium tissue (Table S1). These markers were used to annotate cells with the sctype R package using the SCT scaled data layer. Cells were clustered using the FindNeighbors and FindClusters Seurat functions, setting the resolution parameter to 0.9.

Olfactory gene annotation per cell was performed based on the gene expression levels as calculated with the SCT integration. OR genes with expression level greater than a defined threshold were annotated to the corresponding cell. The threshold was set based on the expression levels of OR genes in the wild type mice, to distinguish the only expressed OR gene from the background noise.

OR genes were mapped to chromosomes based on the mm10 annotation, and the number of chromosomes containing co-expressed OR genes was quantified per cell.

### CUT&RUN data analysis

CUT&RUN data was processed with the nf-core/cutandrun pipeline v3.2.1. In brief, this pipeline performs quality control, adapters trimming, alignment to target genome (GRCm38) and spike-in (D. Melanogaster), reads deduplication, peak calling with MACS2 and consensus peaks detection. MACS2 was used to detect broad peaks. Pipeline parameters were set to: MAPQ > 30 and broad peak cutoff equal to 0.05. From pipeline output gene stack plots were produced. CUT&RUN signal at GI-enhancer coordinates was calculated using the computeMatrix program. The visualization was created with a custom R script. Signal profile was calculated from the signal matrices by summing over columns and smoothing the resulting vector of totals.

### ChIP-seq data analysis

ChIP-seq datasets were first downloaded from the Sequence Reads Archive (SRA) using the nf-core/fetchngs v1.12.0 pipeline. The pipeline downloads and prepares sequencing data from the archive. Subsequently, datasets were processed independently but uniformly with the nf-core/chipseq v2.2.0dev-g671de70 pipeline. ChIP-seq analysis pipeline parameters are summarized in Table S3 and available in the online repository. Briefly, all samples were aligned to the GRCm38 genome assembly and the MGI gene annotation was used. Bowtie2 was used to map sequencing reads to the reference genome. The Encode blacklist file was obtained from online repositories (<u>zenodo.1491733</u>). Broad peak calling was enabled for H3K9me3 samples and, in all cases, Macs2 false discovery ratio threshold was set to 0.05.

H3K9me3 signals from mOSN were compared across OR clusters and promoters of protein coding genes. OR clusters were tiled in 2.2 kb non overlapping windows. For each OR cluster, a window was randomly selected, average signal calculated across the three datasets. Pearson correlation coefficient was then calculated for all combinations of datasets. Asymmetrical windows of 2.2 kb around TSS (2 kb upstream, 200 bp downstream) of randomly selected protein coding genes were generated. Mean signals and correlations were calculated similarly to the OR clusters. Random sampling procedures were repeated 10 times for each feature type. Correlation coefficients were averaged over resampling iterations and converted to distance measurement by subtracting them to 1 (i.e., distance = 1 - average Pearson correlation). Resulting correlation matrices were clustered using hierarchical clustering (with average clustering method) and visualized as dendrograms.

### Hi-C data analysis

Processed Hi-C data was downloaded from the 4D nucleosome project web page. All analyses were performed with the 50 kb resolution. Heatmaps were generated following the method described in Monahan et al. 2019 ^13^. Briefly, the raw matrix was normalized using the Knight-Ruiz method, and bins containing GI-enhancers were subset from genome wide matrices. For downstream analysis, GI-enhancers spanning two bins were discarded (Amorgos).

### Integrated analysis

GI-enhancers were split into two groups by calculating the median CUT&RUN signal value across all replicates (no summarization was performed). GI-enhancers displaying binding signal values above the median were assigned to the high signal group, remainders were assigned to low signal group. The Hi-C signal at GI-enhancers were processed to remove values of enhancers in *cis*, ie. the values corresponding to GI-enhancers on the same chromosome were removed. Total trans interactions were computed by summing all remaining values. The distribution of these values split by CUT&RUN signal was then compared (Figure 2B). OR gene-level annotation about co-expression events was retrieved from the 10X scRNA-seq data. For each OR cluster, the total number of co-expressed genes was calculated. The ratio of co-expressed genes with respect to the total number of genes in the cluster was compared between GI-enhancers split by CUT&RUN signal.

## ACKNOWLEDGMENTS

We thank current and former colleagues from the Boulard lab for their input throughout the project. We acknowledge the critical support of EMBL Rome Core Facilities: Laboratory Animal Facility, Gene Editing and Virus Facility, Light Imaging Facility, GeneCore and PepCore. The authors thank Tim Bestor, Jamie Hacket, Cornelius Gross, and Santiago Rompani for critical reading of the manuscript. This work was supported by open access funding provided by European Molecular Biology. B.A. was supported by a fellowship from the EMBL Interdisciplinary Postdoc (EIPOD) program (Marie Sklodowska-Curie Actions, COFUND grant agreement 847543).

## AUTHOR CONTRIBUTIONS

Conceptualization: MB, FT, BA

Methodology: MB, FT, BA, CL, MA, NH, AS, AHC, EP, MP

Investigation: MB, FT, BA, MM, AR, CL, AS, DG, HA

Visualization: MB, FT, BA, CL, HA, DG, MP

Funding acquisition: MB

Project administration: MB

Supervision: MB

Writing – original draft: MB

Writing – review & editing: MB, HA

## Supplementary Figures Legends

**Figure S1:**
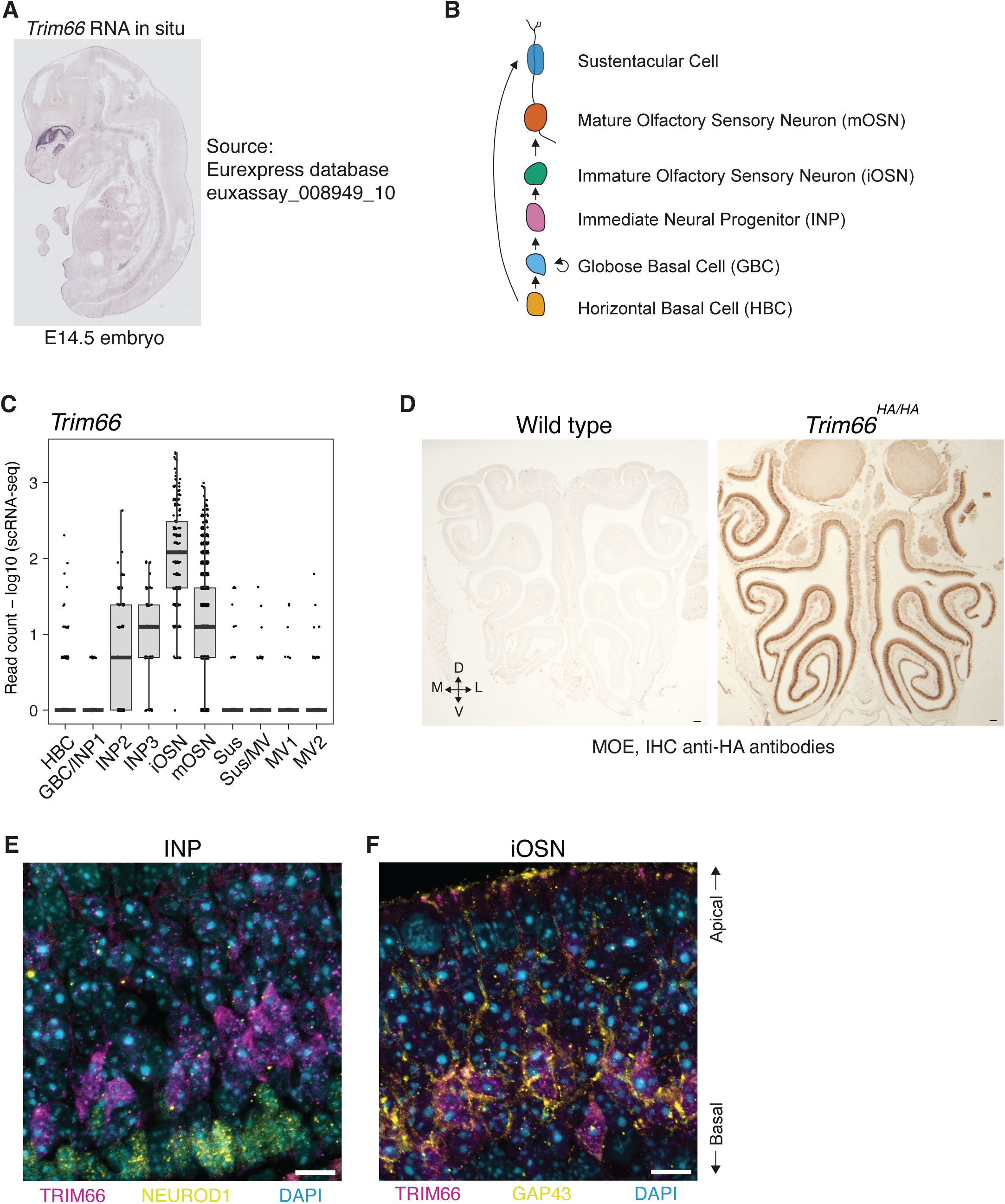
Dynamics of *Trim66* expression during the maturation of olfactory neurons. (A) *In situ* hybridization of *Trim66* mRNA on a sagittal section of WT embryo at E14.5 from the Eurexpress database (Sample ID: euxassay_008949_10) showing a strong and specific signal in the olfactory epithelium ^38^. (B) Schematic representation of the neural lineage differentiation in the main olfactory epithelium. Basal cell layer: horizontal basal cells; apical cell layer: sustentacular cells. (C) scRNA-seq quantification of *Trim66* expression in the main cell types composing the MOE. HBC: horizontal basal cells, GBC: globose basal cells, INP: immediate neural precursors, iOSN: immature olfactory sensory neurons, mOSN: mature olfactory sensory neurons. Sustentacular cell. (D) TRIM66 expression in the adult main olfactory epithelium visualized by immunohistochemistry (IHC). Anti-HA antibodies were used to detect TRIM66-HA on MOE coronal sections from WT (left panel) and *Trim66^HA/HA^* (right panel) animals. The scale bar indicates 100 µm. D: dorsal; L: lateral; M: medial; V: ventral. (E-F) Confocal image of a section of MOE showing TRIM66 expression in INP (E) and iOSN (F). Endogenous TRIM66 fused to the HA epitope tag was detected with anti-HA antibodies (magenta). DNA staining with DAPI is shown in cyan. The scale bar indicates 10 µm. (E) Co-immunostaining of TRIM66-HA (magenta) and NEUROD1 (yellow) as a marker of INP. (F) Co-immunostaining of TRIM66-HA (magenta) and GAP43 (yellow) as a marker of iOSNs.

**Figure S2:**
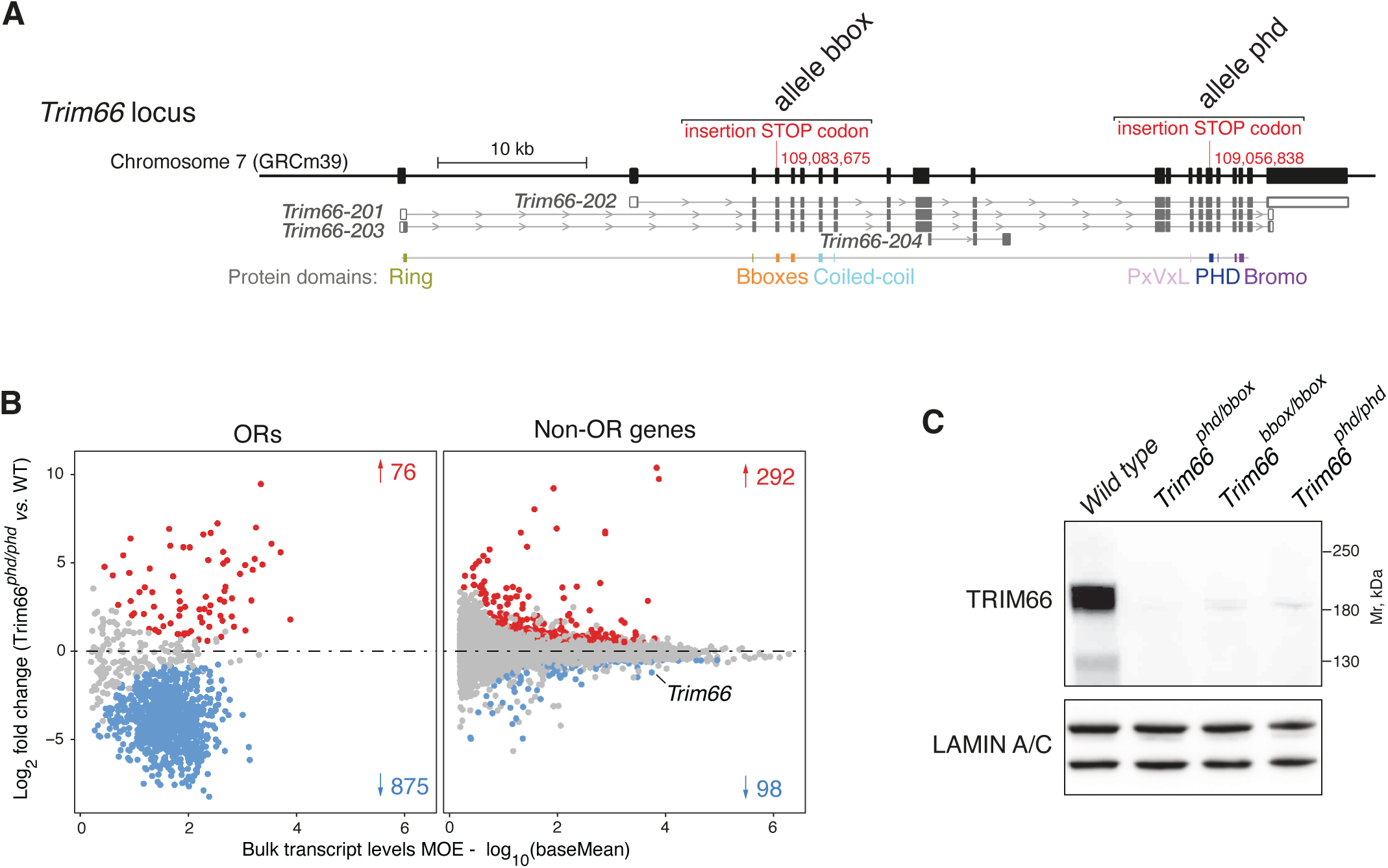
Loss-of-function of *Trim66* deregulates OR genes. (A) Disruption of the murine *Trim66* gene by CRISPR-HDR insertion of a premature STOP codon. Two loss-of-function mutations of *Trim66* were created: *Trim66^bbox^* by insertion of a stop codon in exon 4 (*Trim66-201*); *Trim66^phd^* by insertion of a stop codon in exon 15 (*Trim66-201*). (B) Bulk RNA-seq MA plot depicting differential gene expression in *Trim66^phd/phd^* MOE relative to WT. Significance was defined by a *P*-value lower than 0.01 (Wald test), log2-fold change greater than 0.5 (red dots) or lower than −0.5 (blue dots). (C) Western blot detection of TRIM66 in nuclei isolated from adult MOE of the indicated genotypes. Detection of LAMIN A/C serves as a loading control.

**Figure S3:**
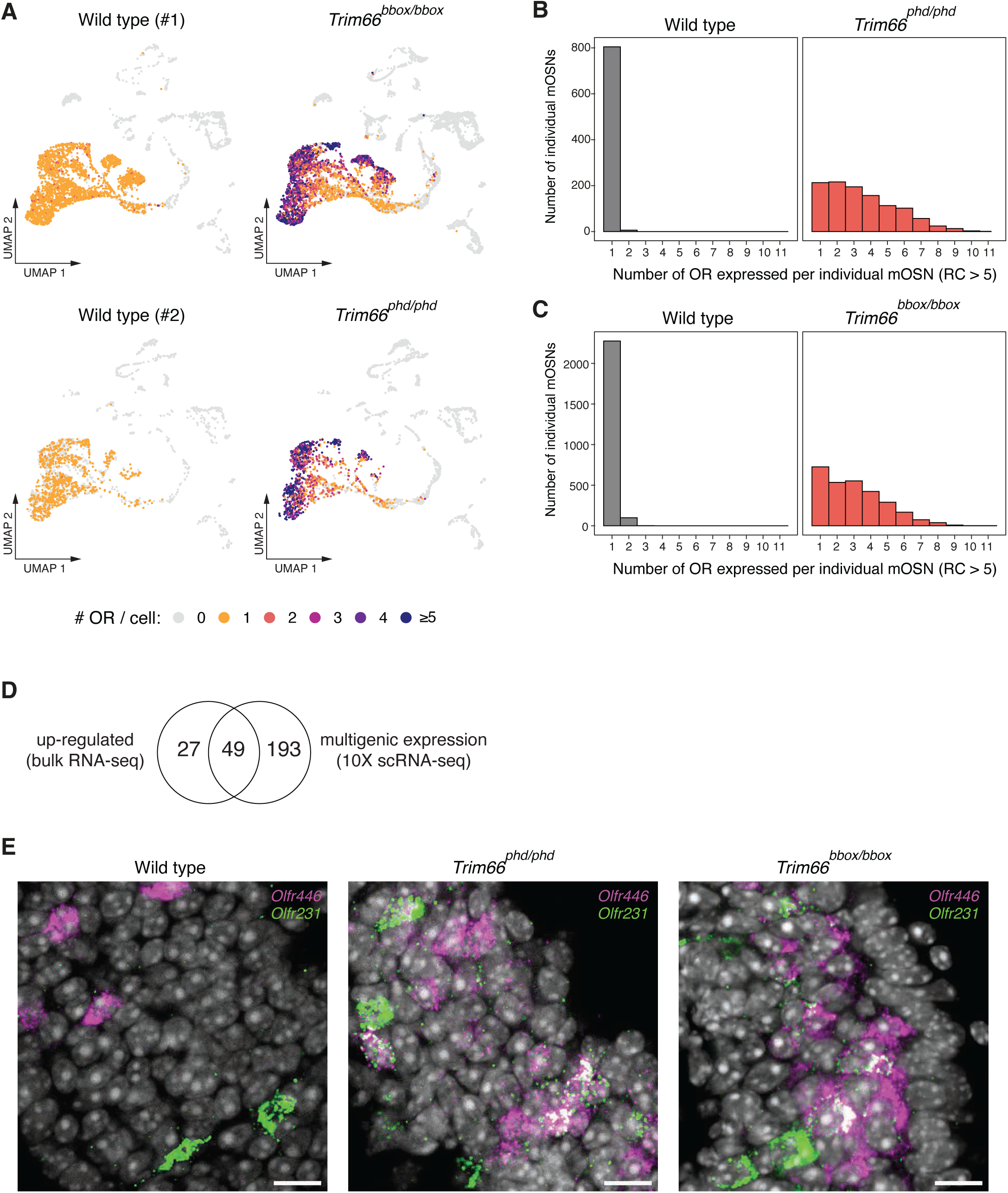
Co-expression of multiple OR genes in *Trim66*-disrupted mOSNs. (A) Uniform manifold approximation and projection (UMAP) of the MOE single-cell RNA-seq data set for the four samples analyzed. The number of different OR mRNAs per single cell is color-coded as indicated. (B) Histogram representing the number of OR genes expressed in individual mOSNs WT (left panel) and *Trim66^phd/phd^*(right panel). Nearly all individual WT mOSNs expressed only one OR gene. *Trim66^phd/phd^*mOSNs expressed a variable number of OR genes ranging from 1 to 11. (C) Histogram representing the number of OR genes expressed in individual mOSNs WT (left panel) and *Trim66^bbox/bbox^*(right panel), reproducing the polygenic expression in the absence of functional *Trim66*. (D) Venn diagram showing common OR genes shared between upregulated genes (bulk RNA-seq dataset) and co-expressed genes (scRNA-seq dataset) in *Trim66^phd/phd^* MOE. (E) Multiplexed Hybridization Chain Reaction (HCR) combined with fluorescence *in situ* hybridization (RNA-FISH) of *Olfr446* (magenta) and *Olfr231* (green) in WT (left panel), *Trim66^phd/phd^* (middle panel), and *Trim66^bbox/bbox^*(right panel) MOE cortical sections. Nuclei were stained by DAPI. The scale bar indicates 10 µm.

**Figure S4:**
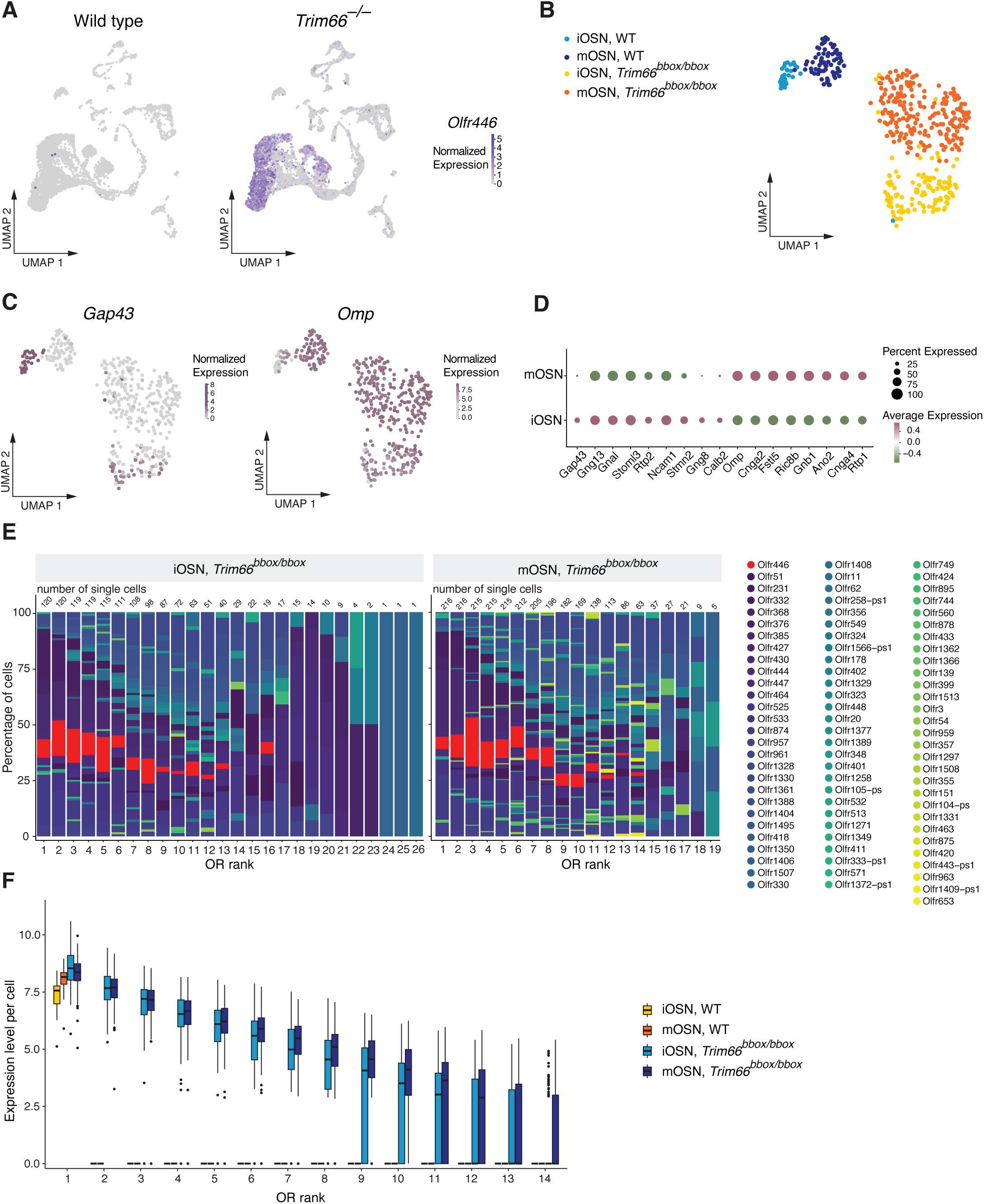
Co-expression of multiple OR genes in *Trim66*-disrupted iOSNs. (A) Uniform Manifold Approximation and Projection (UMAP) of single-cell RNA-seq (10x genomics) integrated two WT MOE (left) and two *Trim66*-deficient MOE (*Trim66^phd/phd^* and *Trim66^bbox/bbox^*; right). The expression of *Olfr446* is shown. (B) UMAP of *Olfr446*+ single-cell RNA-seq (scRNA-seq) dataset sequenced with Smart-seq. Cell type and genotype colors are indicated according to the legend. (C) UMAP visualization of the MOE scRNA-seq of sorted neurons expressing *Olfr446* using the *Olfr446^T2i-TAU-GFP^*reporter allele and sequenced with Smart-seq2. Colors indicate the normalized expression levels of *Gap43* (left) and *Omp* (right). (D) Dotplot showing key marker genes for single neurons *Olfr446+* analyzed by scRNA Smart-seq2. The genes have been selected to annotate the type of each cell. The size of the dot represents the percentage of cells in which the gene is expressed, while the color represents the average expression of the gene within the cell type group. (E) Identity of the co-expressed OR genes in single neurons *Olfr446+* analyzed by scRNA Smart-seq2. For each OR gene rank (x-axis), the stacked bar shows the percentage of cells (y-axis) expressing an OR gene at that expression level. Each color represents a distinct OR gene. Left panel: *Trim66^bbox/bbox^*iOSNs (immature olfactory sensory neurons); right panel: *Trim66^bbox/bbox^* mOSNs (mature olfactory sensory neurons). Numbers above bars indicate the total number of cells analyzed for each rank. (F) OR transcript level in single neurons *Olfr446+* WT and *Trim66^bbox/bbo^*^x^ analyzed by scRNA Smart-seq2. Box plot showing the expression levels of the top 14 OR genes by rank, grouped by cell type and genotype.

